# CIWARS: a web server for waterborne antibiotic resistance surveillance using longitudinal metagenomic data

**DOI:** 10.1101/2024.10.30.621203

**Authors:** Muhit Islam Emon, Yat Fei Cheung, James Stoll, Monjura Afrin Rumi, Connor Brown, Joung Min Choi, Nazifa Ahmed Moumi, Shafayat Ahmed, Haoqiu Song, Justin Sein, Shunyu Yao, Ahmad Khan, Suraj Gupta, Rutwik Kulkarni, Ali Butt, Peter Vikesland, Amy Pruden, Liqing Zhang

## Abstract

The rise of antibiotic resistance (AR) is a major global health crisis, exacerbated by the overuse and misuse of antibiotics, leading to the rapid spread of antibiotic resistance genes (ARGs) in bacterial pathogens. This phenomenon poses significant threats to human and animal health, food security, and economic stability. Water bodies, particularly wastewater treatment plants (WWTPs), serve as critical reservoirs for ARGs, creating environments that favor the proliferation of resistant bacteria. Wastewater-based surveillance (WBS) has emerged as a cost-effective strategy for monitoring AR at the population level, providing real-time data to guide public health and policy decisions. Despite advancements in WBS, there are no comprehensive online analytical platforms for continuous environmental AR surveillance. This paper introduces CIWARS, a web server designed for AR analyses of longitudinal metagenomic data. CIWARS offers comprehensive ARG profiling, taxonomic annotation, and anomalous AR risk points detection. We demonstrate its capabilities through an interactive temporal data visualization, showcasing its potential for enhancing AR risk monitoring and guiding effective mitigation strategies. CIWARS is broadly applicable to longitudinal metagenomic data generated from any environment and aims to support global efforts in addressing the AR crisis by providing cyberinfrastructure for continuous AR surveillance. The web server is freely available at https://ciwars.cs.vt.edu/.

## Introduction

The discovery of antibiotics is considered the cornerstone of modern medicine that saved countless lives from bacterial infection and significantly improved life expectancy [1].

Unfortunately, that cornerstone is now crumbling due to the accelerated spread of antibiotic resistance (AR). While the development of resistance mechanisms in bacteria, referred to as antibiotic resistance genes (ARGs), is a natural evolutionary process, the overuse and misuse of antibiotics have accelerated the emergence and spread of AR in bacteria by creating selection pressure in these environments. AR in bacterial pathogens has now become a worldwide challenge contributing to high morbidity and mortality [2]. The World Health Organization (WHO) named AR among the top 10 threats for global health, and estimated that 10 million deaths could be attributed to AR annually by 2050. If left unchecked, proliferation of drug-resistant pathogens will incur significant costs to both healthcare and economic sectors, threatening human and animal health, and increasing food scarcity and poverty. As such, addressing AR requires concerted efforts at local, national, and global levels.

Although there are numerous avenues through which ARG can disseminate, several studies have highlighted that water bodies, including rivers, lakes, and oceans, as well as wastewater and drinking water systems, serve as reservoirs and conduits for ARGs [3-8].

Wastewater treatment plants (WWTPs) are especially regarded as major reservoirs of resistance genes because they receive wastewater from various sources, including households, hospitals, industries, and agricultural runoff which are often polluted with large amounts of antibiotics, antibiotic-resistant bacteria (ARB), and their resistance genes. The presence of antibiotics in the wastewater creates a selection pressure that favors the survival and proliferation of resistant bacteria by facilitating the horizontal gene transfer of resistance genes.

Inevitably, Wastewater based surveillance (WBS) has emerged as the most cost-effective approach for predicting the occurrence of ARB at the population level [9-16]. Monitoring antimicrobial resistance (AMR) in wastewater is useful in understanding the One Health approach, which recognizes the interconnectedness of human, animal, and environmental health and the presence of AMR as a contaminant found throughout [17].

Furthermore, WBS provides an opportunity to detect AMR trends in real-time or even develop an early prediction tool for future infection outbreaks caused by ARB [18]. Such information is essential to guide healthcare and public policymaking in developing cost-effective strategies for mitigating challenges associated with AMR.

Much effort has been made to develop and implement wastewater surveillance systems for AMR. Notably, the CDC recently announced efforts to expand the National Wastewater Surveillance System to include environmental AMR. As a result, a substantial amount of metagenomic data is being generated, with a notable increase in longitudinal data. However, there is still no online comprehensive environmental AR surveillance system that facilitates monitoring, evaluation, and reporting of AR risk indicators. In this paper, we introduce the CyberInfrastructure for Waterborne Antibiotic Resistance Risk Surveillance (CIWARS), a web server designed to facilitate longitudinal metagenomic data analyses through read matching and assembly-based annotation pipelines. In addition to comprehensive ARG profiling and taxonomic annotation, CIWARS also offers detection of anomalous time points in terms of AR risks. CIWARS can be applied to longitudinal metagenomic samples from any environment. The web service is publicly available at https://ciwars.cs.vt.edu/.

## Materials and Method

Raw paired-end metagenomic short-read data is submitted to the CIWARS server through a user-friendly web interface. Users are required to create an account and a profile on the server. This profile enables them to submit, analyze, retrieve, and compare AR risk across their longitudinal samples. CIWARS organizes metagenomic samples and corresponding analysis results into user-specific projects.

### Required data types

CIWARS requires users to upload raw paired-end short-read sequences in gzipped FASTQ format. Additionally, users must submit a metadata file (format provided by the server) that includes the sequencing time for each metagenomic sample. Other sample metadata, such as pH, temperature, or any additional information of interest to the user, are optional.

### Analysis pipelines

After being stored on the CIWARS server, raw reads are queued for taxonomic and ARG annotations, along with AR risk assessments. CIWARS utilizes two distinct pipelines: the read-matching pipeline and the assembly-based pipeline (Fig. 1).

**Figure 1.**
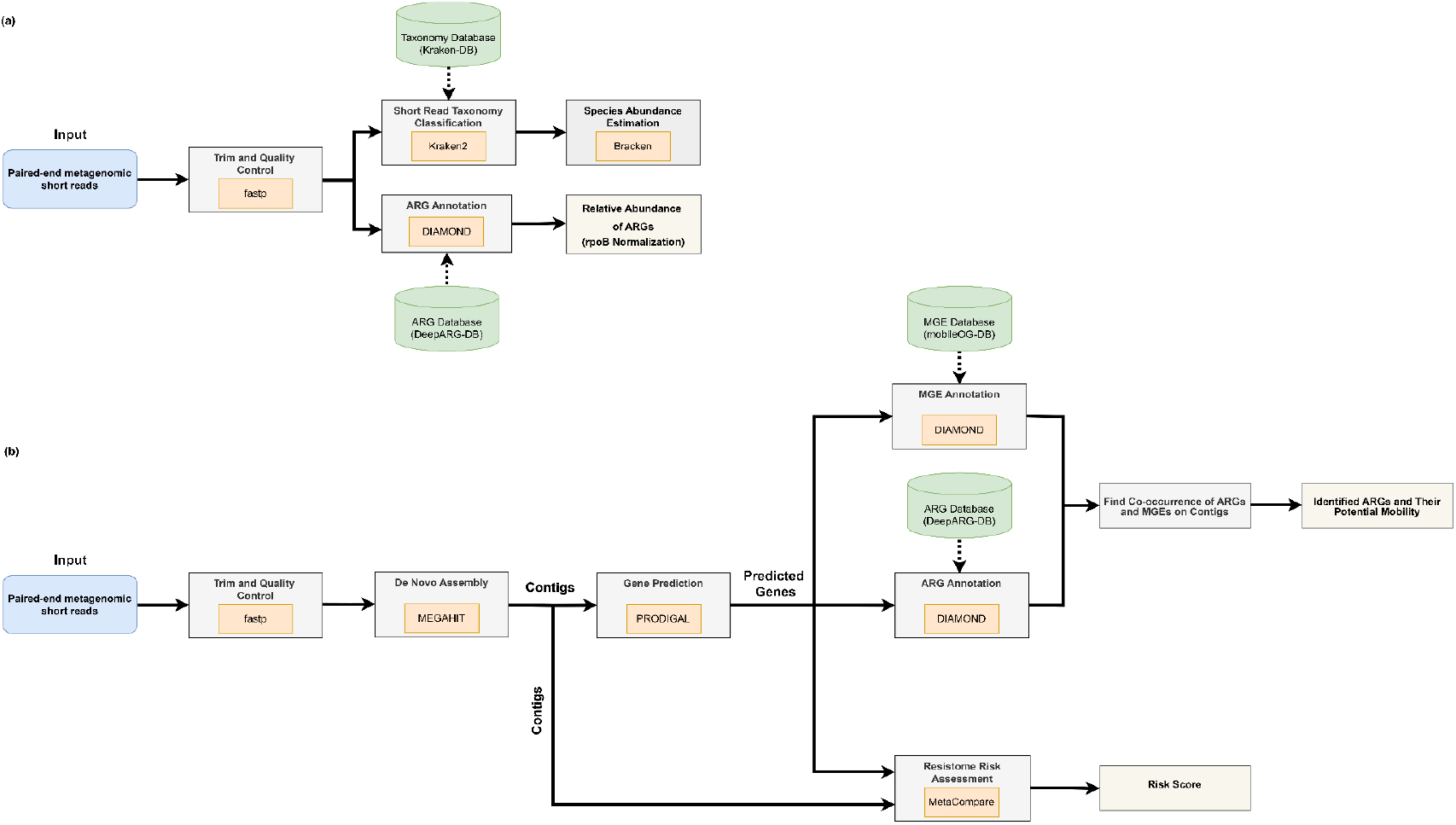
Overview of the computational pipelines implemented in the CIWARS service for taxonomic and ARG annotation, as well as AR risk assessment. (a) Read matching pipeline. (b) Assembly pipeline

#### Read matching pipeline

The read matching pipeline conducts taxonomic and ARG annotation of metagenomic data comparing the raw sequence reads to reference databases. Initially, the reads are trimmed and quality-filtered using fastp [19] with options (--trim_poly_g, --detect_adapter_for_pe, --trim_poly_x, --average_qual 10) to remove low-quality sequences.

For taxonomic annotation, Kraken2 [20] is employed, utilizing its Standard-16 taxonomy database, which is capped at 16 GB, for the rapid classification of quality-controlled metagenomic reads down to the species level. Bracken [21] is then used to estimate relative abundances of species based on Kraken2 classification.

Following filtering and trimming, paired-end reads are merged by aligning overlapping regions using VSEARCH (22). This merging process generates longer reads, which can enhance the performance of read mapping algorithms. Next, ARG annotation is performed by aligning reads against the ARG reference database DeepARG-DB [23] using the DIAMOND BLASTX aligner [24]. The representative hit approach is applied with alignment criteria set to e-value < 1e-10 and identity > 80%. To facilitate comparison across time points, we calculate the relative abundance of ARGs, enabling users to visualize similarities and differences among samples. A key component of this comparison process is data normalization. We employ the “ARGs/cell” normalization approach which is more meaningful from a biological standpoint [25]. We estimate the ARGs/cell metric by dividing the number of ARGs by the single-copy gene rpoB (the β subunit of bacterial RNA polymerase), as outlined by Thornton et al. [26] and Zhang et al. [27].

#### Assembly pipeline

In the assembly process, metagenomic reads are merged into long contiguous sequences, yielding richer genomic information than raw reads. Like the read-matching pipeline, input reads are first cleaned with fastp. MEGAHIT [28], with the “--meta-sensitive” option, is then used to assemble these quality-controlled, trimmed paired-end reads into contigs. Once contigs are assembled, PRODIGAL [29] (-p meta), a microbial gene prediction tool, identifies genes within each contig.

To compute the AR risk score for the input sample, we employ MetaCompare [30], a computational framework that takes contigs and predicted genes as inputs to evaluate the resistome risk from metagenomic sequencing data. This metric is calculated based on the estimated co-occurrence of ARGs, mobile genetic elements (MGEs), and pathogens on the same contig.

For ARG annotation, the predicted genes (translated proteins from PRODIGAL) are compared to the ARG reference database DeepARG-DB using the DIAMOND BLASTP aligner. We employ the best hit strategy with an e-value < 1e-10, identity > 60%, and coverage > 70%. Additionally, we assess the potential mobility of ARGs by determining whether they are associated with MGEs. Specifically, an ARG is considered to exhibit mobility if it is carried by an MGE. To identify contigs derived from MGEs, we align the predicted genes against the mobile orthologous groups database (mobileOG-DB) [31] using the DIAMOND BLASTP aligner, again employing the best hit strategy with an e-value < 1e-10, identity > 60%, and coverage > 70%. If an ARG co-occurs with a gene matched in mobileOG-DB on the same contig, we classify the contig as potentially derived from an MGE and annotate the ARG as having potential mobility.

### Anomaly detection

Interquartile range (IQR)-based statistical method is implemented in the server to identify the anomaly candidates. IQR is calculated as the range between the first (Q1) and the third (Q3) quartiles of the dataset and the lower and upper bounds of the dataset are defined by Q1-1.5^*^IQR and Q3+1.5^*^IQR, respectively. Any data point that lies below the lower bound or above the upper bound is considered an anomaly candidate.

### Visualization of time series results

Users can visualize the ARG profiles and taxonomy abundances over time in stacked bar plots and the detailed percentages of the selected time point will be shown in the piechart. Heatmaps are also produced for Pearson correlation between Operational Taxonomic Units (OTUs) in the user-selected taxonomy level and also correlations between the timepoints. For anomaly visualization, users can select different targets to examine their pattern changes over time.

Anomalies will be marked and the number of anomalies at specified times will be counted for inferences of anomalous samples at certain time points.

### Web server and system architecture

The overall system architecture of CIWARS is shown in Fig. 2. CIWARS is built on a layered, cloud-based architecture, utilizing Docker containers running under Kubernetes, deployed via Helm on AWS resources. While targeting the AWS environment, few parts of the application are tied directly to AWS, and it should be feasible to host on other cloud platforms with minor modifications. The layers of the application include: Authentication, Database, REST API, Front End, Queue, Pipeline Orchestration, and Filesystem.

**Figure 2.**
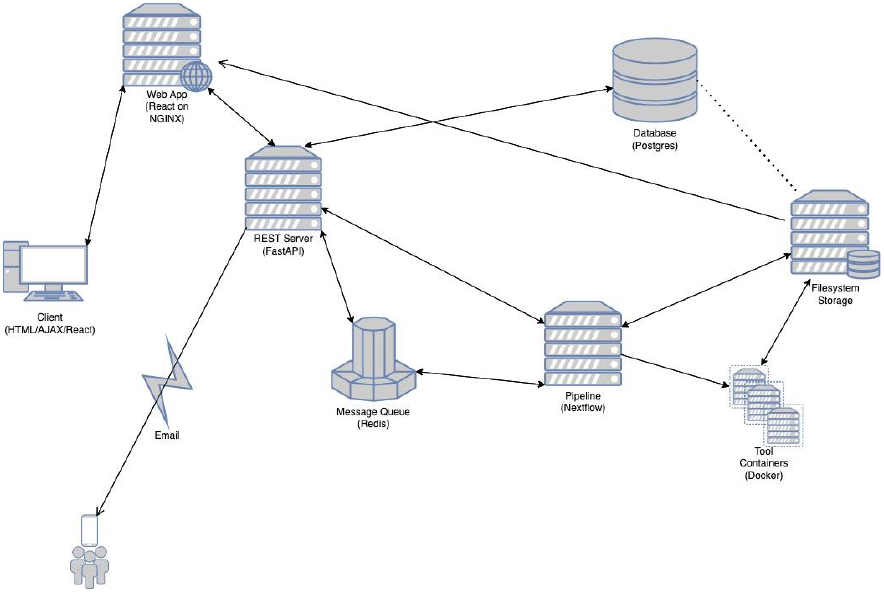
The system architecture of CIWARS.

Authentication is required for use of CIWARS, and authentication is implemented via the OAUTH2 protocol using the /authenticate scope. The initial implementation integrates ORCID as an OAUTH provider. Additional providers can be easily integrated as needed.

Underlying the application, the data layer is implemented via a Postgres database. Application data and metadata are stored in this database, and access to the database is enabled through a REST API using the Tortoise ORM framework for database interaction. Aerich is used for migration/versioning of the database schema.

The REST API is implemented in Python, using the FastAPI REST framework, using the Pydantic framework for message structure and validation. The REST API, in concert with the Authentication layer, makes use of encrypted JWT tokens for authorization control, ensuring that users can only access their own data. All interactions between layers take place through the REST API.

The Queue employs Redis and Celery for asynchronous task management, running jobs on AWS Spot instances to save costs. Interrupted tasks are re-enqueued with higher priority, leveraging Nextflow’s reentrancy capabilities to resume from the interruption point. Pipeline Orchestration uses NextFlow and Docker for multi-step processes, dynamically spawning Kubernetes Jobs on Spot instances. New pipelines can be added through Docker containers without altering the system’s codebase.

The Filesystem provides storage for user-provided sample input files, Nextflow working files and logs (the means by which Nextflow provides reentrancy), pipeline output files, and visualization output files for pipelines that provide visualizations. There are user quotas to limit the number of concurrent input and output files in use by an individual user at any given time, in order to keep storage needs within reasonable limits.

## Example Analyses

### Antimicrobial resistance profiling in WWTP samples

We conducted an in-depth AMR profiling study on 15 samples collected from a WWTP in Christiansburg, Virginia, comprising both influent and effluent wastewater. The analysis is detailed through nine figures, each providing insights into temporal changes in AR.

### ARG drug class abundances over time

Fig. 3(a) depicts the abundance of ARG drug classes at specific time points and their temporal changes. The stacked bar plot visually represents the abundance of ARGs across different drug classes at different time points. Clicking on a specific stack within the bar plot triggers a pie chart display, providing a detailed breakdown of the ARGs associated with the chosen drug class for that time. Hovering over the pie chart segments reveals the exact abundance values for the selected time point (Fig. 3(b)). Users can reveal the periods of increase or decrease in the abundance of different ARG drug classes and determine which ARG drug classes are most prevalent at specific time points. From the figure, they can also detect the emergence or disappearance of specific ARG drug classes, which can indicate changes in antibiotic resistance patterns.

**Figure 3.**
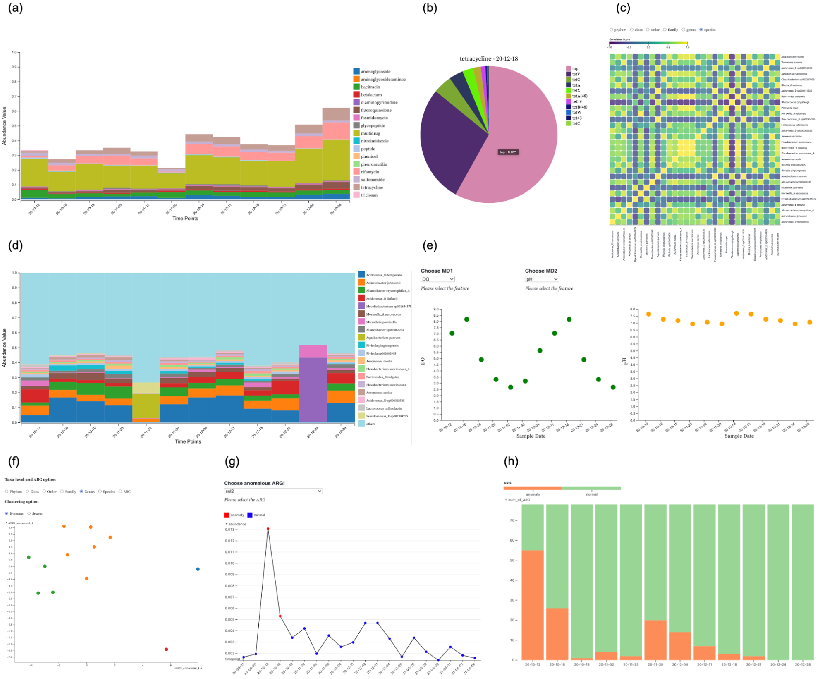
Visualization provided in CIWARS. (a) ARG drug class abundances over time. (b) Pie chart showing the percentage of each ARG in the selected drug class for the selected time point. (c) OTU correlation heatmap. (d) Bacterial class abundances over time. (e) Metadata features trends over time. (f) Scatterplot for the clustering results based on the selected taxa-level abundance. (g) Anomalous time points for ARGs. (h) Summary of the anomalous ARGs across the time points.

### OTU correlation and time point correlation heatmap

The heatmap visualizes the degree of correlation between OTUs across all samples (Fig. 3(c)). To ensure easy understanding, the top 30 OTUs with the highest relative abundance will be selected for visualization and Pearson correlation will be measured using all the longitudinal samples. Correlation heatmap will be plotted for each taxa level including phylum, class, order, family, genus, and species. This visualization will help the user understand the correlation levels and relationships between OTUs at each taxa level over the time points. We also provide the heatmap showing the degree of Pearson correlation between time points based on the relative abundance of all species, allowing the user to check for correlations between time points, months, or seasons.

### Bacterial abundances over time

Fig. 3(d) illustrates the abundance of bacteria at specific time points and how their composition changes over time. The stacked bar charts display the taxa-level abundances at various time points. By selecting a taxa level using radio buttons, users can view bacterial abundance values for the chosen level. We retained only the top 20 dominant entities in the time-series dataset for the selected taxa level. This figure will help users identify which bacterial taxa are dominant at specific time points and how their dominance shifts. Users can also assess the stability or variability of the bacterial community over time, determining periods of relative stability or significant change.

### Metadata feature trends over time

Fig. 3(e) visualizes trends in metadata features over time. Users can select two features to visualize simultaneously, and hovering over each point highlights the corresponding value of the other feature at the same time point in red. This visualization helps users examine changes in the selected features and determine whether the two features might be associated.

### Scatterplot for the clustering results based on the selected taxa-level abundance

The scatterplot in Fig. 3(f) displays samples (time points) based on the abundances for the selected taxa level. Time points are clustered based on ARG distribution using non-metric multidimensional scaling (NMDS) and K-means clustering, with the value of K selected based on the silhouette score. Users can view clustered time points by K-means or by season (spring, summer, autumn, winter). Hovering over the points reveals the specific time points.

### Anomalous time points for ARGs and metadata features

We identify anomalous time points for each ARG by showing the trend of rpoB-normalized abundance for user-selected ARGs across different time points, with anomalous time points highlighted in red (Fig. 3(g)). The anomalies were identified based on IQR rules. This visualization can serve as an early warning system for potential surges in ARG abundances or provide insights into potential disease outbreaks derived from ARG data. In addition to identifying anomalous time points for ARGs, we also detect anomalous time points for each metadata feature provided by the user and display the trends of metadata feature value changes. Anomalies, identified using IQR rules, are highlighted in red. By comparing the visualizations of anomalous time points identified by either ARG abundance or metadata features, the user can determine if changes in ARG abundance may be influenced by metadata factors such as experimental conditions or sampling.

### Summary of the anomalous ARGs across the time points

To provide an overview of the number of potential anomalous ARGs and how they change over time, a summary of the ARG anomalies identified by IQR rules and the normal ones for each sample is presented (Fig. 3(h)). This summary will help users examine the temporal changes in the number of anomalies, identifying periods of increased or decreased ARG activity.

Additionally, understanding the frequency of anomalies can help assess the stability or variability of ARG levels within the samples, prompting further investigation into sample collection or processing methods.

The plots shown in Fig. 3 collectively provide a comprehensive view of the AR landscape in the WWTP samples, highlighting temporal dynamics and potential anomalies.

## REFERENCES

1. Caracciolo AB, Visca A, Massini G, Patrolecco L, Miritana VM, Grenni P. Environmental fate of antibiotics and resistance genes in livestock waste and digestate from biogas plants. World Health. 2020;21:23

2. Akova M. Epidemiology of antimicrobial resistance in bloodstream infections. Virulence. 2016 Apr 2;7(3):252–66

3. Goulas A, Belhadi D, Descamps A, Andremont A, Benoit P, Courtois S, Dagot C, Grall N, Makowski D, Nazaret S, Nélieu S. How effective are strategies to control the dissemination of antibiotic resistance in the environment? A systematic review. Environmental Evidence. 2020 Dec;9:1–32

4. Li, S., Ondon, B.S., Ho, S.H., Zhou, Q. and Li, F., 2023. Drinking water sources as hotspots of antibiotic-resistant bacteria (ARB) and antibiotic resistance genes (ARGs): Occurrence, spread, and mitigation strategies. Journal of Water Process Engineering, 53, p.103907

5. Nnadozie CF, Odume ON. Freshwater environments as reservoirs of antibiotic resistant bacteria and their role in the dissemination of antibiotic resistance genes. Environmental pollution. 2019 Nov 1;254:113067

6. Suzuki S, Pruden A, Virta M, Zhang T. Antibiotic resistance in aquatic systems. Frontiers in microbiology. 2017 Jan 25;8:14

7. Wellington EM, Boxall AB, Cross P, Feil EJ, Gaze WH, Hawkey PM, Johnson-Rollings AS, Jones DL, Lee NM, Otten W, Thomas CM. The role of the natural environment in the emergence of antibiotic resistance in Gram-negative bacteria. The Lancet infectious diseases. 2013 Feb 1;13(2):155–65

8. Karkman A, Do TT, Walsh F, Virta MP. Antibiotic-resistance genes in waste water. Trends in microbiology. 2018 Mar 1;26(3):220–8

9. Ng C, Tay M, Tan B, Le TH, Haller L, Chen H, Koh TH, Barkham TM, Thompson JR, Gin KY. Characterization of metagenomes in urban aquatic compartments reveals high prevalence of clinically relevant antibiotic resistance genes in wastewaters. Frontiers in microbiology. 2017 Nov 16;8:2200

10. Khan FA, Hellmark B, Ehricht R, Söderquist B, Jass J. Related carbapenemase-producing Klebsiella isolates detected in both a hospital and associated aquatic environment in Sweden. European Journal of Clinical Microbiology & Infectious Diseases. 2018 Dec;37:2241–51

11. Hutinel M, Huijbers PM, Fick J, Åhrén C, Larsson DG, Flach CF. Population-level surveillance of antibiotic resistance in Escherichia coli through sewage analysis. Eurosurveillance. 2019 Sep 12;24(37):1800497

12. Pärnänen KM, Narciso-da-Rocha C, Kneis D, Berendonk TU, Cacace D, Do TT, Elpers C, Fatta-Kassinos D, Henriques I, Jaeger T, Karkman A. Antibiotic resistance in European wastewater treatment plants mirrors the pattern of clinical antibiotic resistance prevalence. Science advances. 2019 Mar 27;5(3):eaau9124

13. Aarestrup FM, Woolhouse ME. Using sewage for surveillance of antimicrobial resistance. Science. 2020 Feb 7;367(6478):630–2

14. Huijbers PM, Larsson DJ, Flach CF. Surveillance of antibiotic resistant Escherichia coli in human populations through urban wastewater in ten European countries. Environmental Pollution. 2020 Jun 1;261:114200

15. Sims N, Kasprzyk-Hordern B. Future perspectives of wastewater-based epidemiology: monitoring infectious disease spread and resistance to the community level. Environment international. 2020 Jun 1;139:105689

16. Blaak H, Kemper MA, de Man H, van Leuken JP, Schijven JF, van Passel MW, Schmitt H, de Roda Husman AM. Nationwide surveillance reveals frequent detection of carbapenemase-producing Enterobacterales in Dutch municipal wastewater. Science of The Total Environment. 2021 Jul 1;776:145925

17. Tiwari A, Kurittu P, Al-Mustapha AI, Heljanko V, Johansson V, Thakali O, Mishra SK, Lehto KM, Lipponen A, Oikarinen S, Pitkänen T. Wastewater surveillance of antibiotic-resistant bacterial pathogens: A systematic review. Frontiers in microbiology. 2022 Dec 15;13:977106

18. Flach CF, Hutinel M, Razavi M, Åhrén C, Larsson DJ. Monitoring of hospital sewage shows both promise and limitations as an early-warning system for carbapenemase-producing Enterobacterales in a low-prevalence setting. Water research. 2021 Jul 15;200:117261

19. Chen S, Zhou Y, Chen Y, Gu J. fastp: an ultra-fast all-in-one FASTQ preprocessor. Bioinformatics. 2018 Sep 1;34(17):i884–90

20. Wood DE, Lu J, Langmead B. Improved metagenomic analysis with Kraken 2. Genome biology. 2019 Dec;20:1–3

21. Lu J, Breitwieser FP, Thielen P, Salzberg SL. Bracken: estimating species abundance in metagenomics data. PeerJ Computer Science. 2017 Jan 2;3:e104

22. Rognes T, Flouri T, Nichols B, Quince C, Mahé F. VSEARCH: a versatile open source tool for metagenomics. PeerJ. 2016 Oct 18;4:e2584

23. Arango-Argoty G, Garner E, Pruden A, Heath LS, Vikesland P, Zhang L. DeepARG: a deep learning approach for predicting antibiotic resistance genes from metagenomic data. Microbiome. 2018 Dec;6:1–5

24. Buchfink B, Reuter K, Drost HG. Sensitive protein alignments at tree-of-life scale using DIAMOND. Nature methods. 2021 Apr;18(4):366–8

25. Davis BC, Brown C, Gupta S, Calarco J, Liguori K, Milligan E, Harwood VJ, Pruden A, Keenum I. Recommendations for the use of metagenomics for routine monitoring of antibiotic resistance in wastewater and impacted aquatic environments. Critical Reviews in Environmental Science and Technology. 2023 Oct 2;53(19):1731–56

26. Thornton CN, Tanner WD, VanDerslice JA, Brazelton WJ. Localized effect of treated wastewater effluent on the resistome of an urban watershed. GigaScience. 2020 Nov;9(11):giaa125

27. Zhang SY, Tsementzi D, Hatt JK, Bivins A, Khelurkar N, Brown J, Tripathi SN, Konstantinidis KT. Intensive allochthonous inputs along the Ganges River and their effect on microbial community composition and dynamics. Environmental microbiology. 2019 Jan;21(1):182–96

28. Li D, Liu CM, Luo R, Sadakane K, Lam TW. MEGAHIT: an ultra-fast single-node solution for large and complex metagenomics assembly via succinct de Bruijn graph. Bioinformatics. 2015 May 15;31(10):1674–6

29. Hyatt D, Chen GL, LoCascio PF, Land ML, Larimer FW, Hauser LJ. Prodigal: prokaryotic gene recognition and translation initiation site identification. BMC bioinformatics. 2010 Dec;11:1–1

30. Oh M, Pruden A, Chen C, Heath LS, Xia K, Zhang L. MetaCompare: a computational pipeline for prioritizing environmental resistome risk. FEMS microbiology ecology. 2018 Jul;94(7):fiy079

31. Brown CL, Mullet J, Hindi F, Stoll JE, Gupta S, Choi M, Keenum I, Vikesland P, Pruden A, Zhang L. mobileOG-db: a manually curated database of protein families mediating the life cycle of bacterial mobile genetic elements. Applied and environmental microbiology. 2022 Sep 22;88(18):e00991–22

